# Collagen staining with fast green FCF enables 3D imaging of pulmonary fibrosis

**DOI:** 10.64898/2026.08.27.747478

**Authors:** Mahhum Saqib, Amy K. Rivers, Sylvia Masala, Jonathan R. Baker, Carl Hobbs, Andreas Bodén, Ani Augustine Jose, Dylan Herzog, Simon J. Cleary

**Affiliations:** Institute of Pharmaceutical Science and King’s Centre for Lung Health, King’s College London, London, UK; School of Immunology & Microbial Sciences and King’s Centre for Lung Health, King’s College London, London, UK; Wolfson Sensory, Pain and Regeneration Centre, King’s College London, London, UK; Microscopy Innovation Centre, King’s College London, London, UK

**Author notes:** Co-first authors (Mahhum Saqib, Amy K. Rivers).

## Abstract

Current approaches for imaging fibrotic remodeling have sensitivity, specificity and cost drawbacks that limit both preclinical research and clinical diagnosis. Here, we show that fast green FCF, a small molecule that binds to fibrillar collagen, enables highly sensitive and specific imaging of fibrosis in lung samples from mice and humans using fluorescence microscopy. We report strategies for using fast green FCF staining to assess fibrotic remodeling using precision-cut lung slice and whole-biopsy preparations. Our findings demonstrate that fluorescence imaging of fast green FCF-stained collagen will be useful for fibrosis research and may help to improve detection of fibrosis in clinical pathology.

## Introduction

Fibrosis, the pathological replacement of healthy tissue with non-functional scar tissue, leads to substantial morbidity and mortality. Dysregulated fibrosis within the lungs becomes life-threatening in idiopathic pulmonary fibrosis (IPF) and in several other forms of interstitial lung disease that cause progressive pulmonary fibrosis (PPF). IPF and PPF are incurable and typically progress towards end-stage lung failure within 3-5 years of diagnosis^1,2^.

Key priorities for research aimed at improving pulmonary fibrosis outcomes include developing improved treatments and enabling early detection^3^. Unfortunately, both preclinical research and early diagnosis are currently made challenging by limitations of current approaches for measuring and detecting fibrosis. In experimental studies and in clinical pathology, fibrosis is assessed using formalin-fixed paraffin-embedded lung tissue samples that are sectioned and stained with dyes that label collagen^2,4^. Quantification usually involves manual scoring of fibrotic pathology on sample fields^5^, a process that is time-consuming, can vary between assessors and can under-sample pathology. Immunofluorescence staining of collagen can be useful but relies on expensive antibodies that sometimes incompletely penetrate dense fibrotic tissue, which can result in false negative or artifactually dim staining^6–8^. Radiological imaging methods are used in clinical and experimental fibrosis studies but currently lack the resolution and specificity needed to differentiate fibrotic changes from inflammation^9^. Total lung collagen can be estimated from tissue homogenates, but these assays produce datasets with high variance and can fail to detect resolution of fibrosis^4,10^.

Fast green FCF, a dye that binds collagen and has been used in variants of Masson’s trichrome staining, has recently been identified as a fluorophore useful for fluorescence microscopy^7^. Fast green FCF is an inexpensive small molecule that readily diffuses through dense tissue structures, staining collagen with remarkable brightness and specificity^7^. Because of these features, we hypothesized that fast green FCF staining might be useful help improve assessment of pulmonary fibrosis pathology.

## Results

Adapting strategies that have been used for preparations including mouse embryos and human skin^7,11^, we developed protocols for staining collagen in mouse precision-cut lung slices with fast green FCF (**Fig 1A**). Excitingly, this approach revealed the type I collagen structures that structurally support lung alveoli with unprecedented detail and preservation of morphology (**Fig 1B,C**)^12^. The non-fibrillar type IV collagen supporting capillary basement membranes was not labelled^8,12^, indicating specificity of fast green FCF for fibrillar collagen. As excitation-emission spectra for fast green FCF have not been published, we used lambda scans to determine that collagen-bound fast green FCF is maximally excited at 630 nm with peak emission at 663 nm (**Fig 1D**).

**Figure 1.**
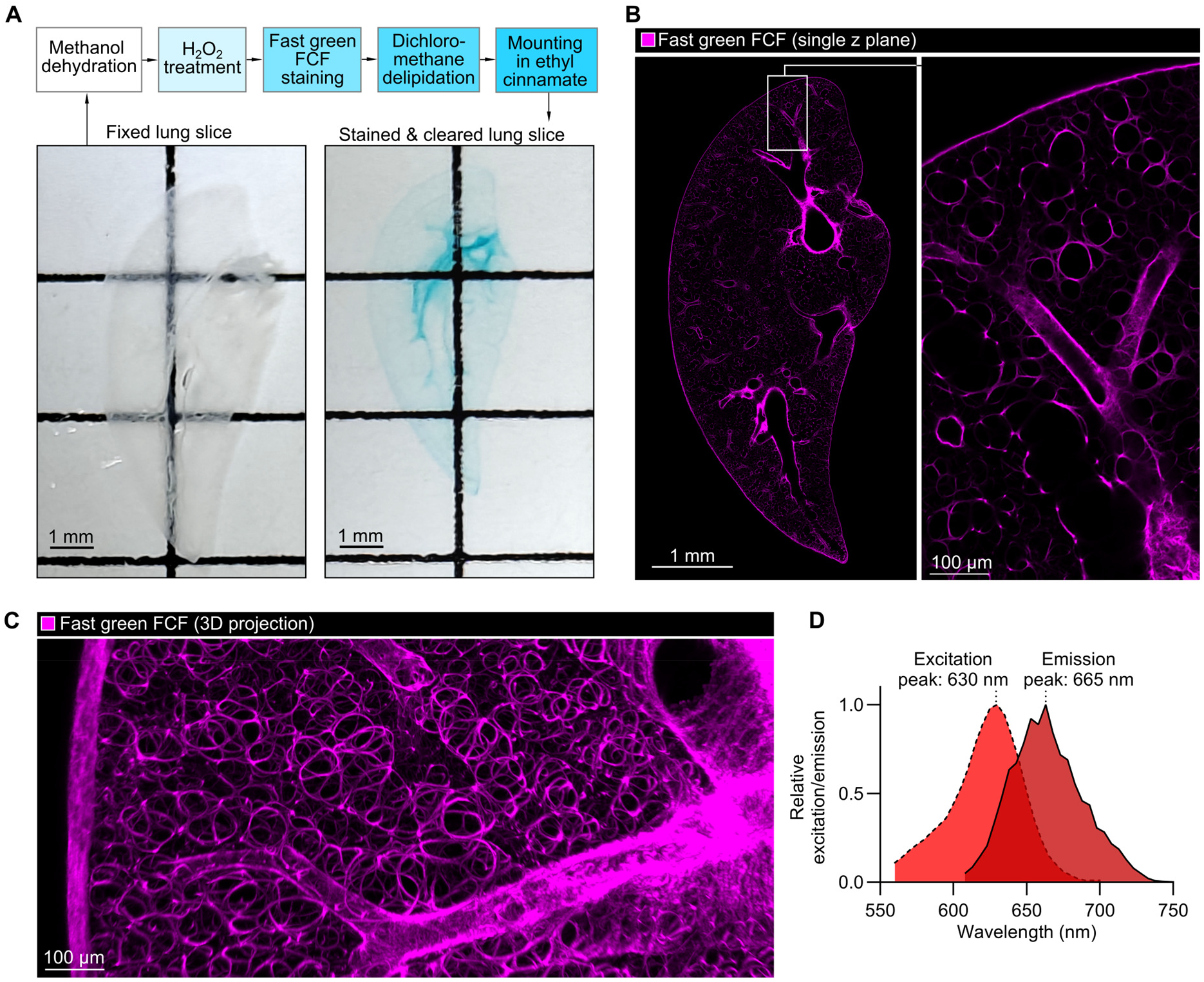
Fast green FCF staining in cleared precision-cut lung slices reveals the delicate collagen network in lung parenchyma. **A**. Representative images of precision-cut mouse lung slices before and after staining and clearing, with summary of workflow. **B**. Image of a single z plane of a precision-cut lung slice stained with fast green FCF. **C**. 3D projection (86 µm thickness, 2 µm z slice increments) from the same sample showing collagen structure in lung parenchyma. **D**. Excitation/emission profile of fast green FCF in ethyl cinnamate mountant.

To test the suitability of fast green FCF for imaging fibrotic changes, we stained lung slices from healthy control mice and from littermates treated with bleomycin, a chemotherapeutic that causes and is widely used to model pulmonary fibrosis. Lungs were harvested 2 weeks after dosing, when inflammatory injury has occurred and fibrotic changes are developing in bleomycin-treated mice^6^. Pathological changes were visible in lungs from bleomycin-treated mice from fast green FCF signal alone (**Fig 2A**). These changes could be quantified as simple increases in fast green FCF signal intensity in lung parenchyma and increases in the percentage of lung parenchyma positive for fast green FCF (**Fig 2B,C**). Notably, the collagen structures that fast green FCF clearly outlines in the healthy alveolar interstitium are less completely resolved with use of the industry standard Masson’s trichrome staining method due to limitations of structural preservation in thin paraffin sections and the challenge of deconvolution of signal from different stains in brightfield microscopy (**Fig 2D**). Picrosirius red staining reveals collagen fibrils in the alveolar interstitium and emits fluorescence but also labels other structures (**Fig 2E,F**). This lack of specificity was visible in mixed inflammatory-fibrotic lesions in lungs of bleomycin-treated mice, in which picrosirius red staining and fluorescence highlighted cytoplasm of some cells as well as non-fibrillar basement membrane collagen fibers (**Fig 2E,F**). The specificity of picrosirius red for fibrillar collagen can be improved by looking for birefringence using polarized light^13^, but imaging birefringence is not well-suited for quantification or 3D imaging. In contrast to traditional histology approaches, fast green FCF staining was compatible with 3D imaging in fibrotic samples (**Fig 3A**). Therefore, fast green FCF staining of precision-cut lung slices has advantages over traditional histology methods of enabling more quantitative, complete and specific visualization of the collagen network in lung parenchyma.

**Figure 2.**
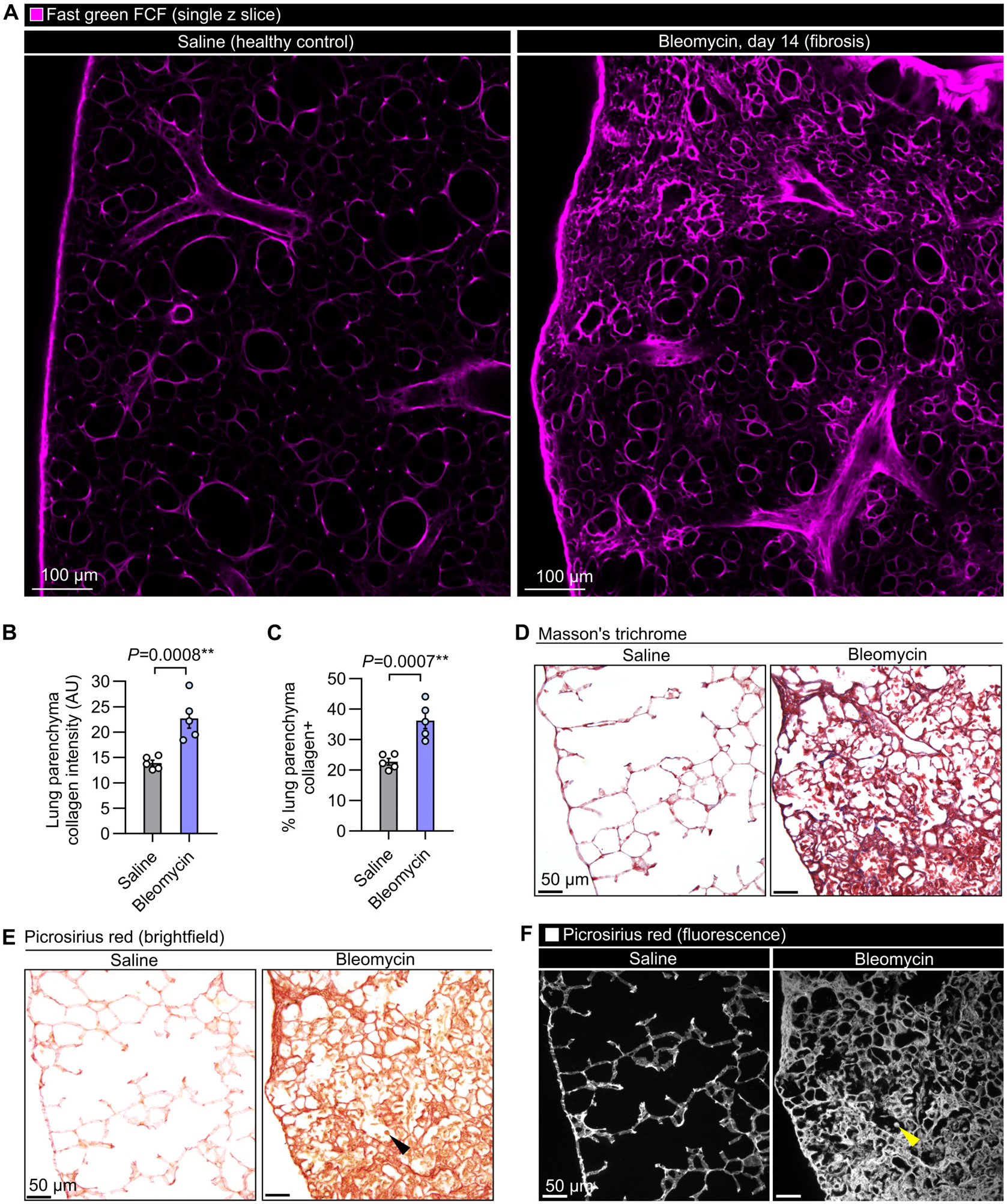
Fast green FCF staining enables more complete and specific quantification of increases in collagen deposition than established histological approaches. **A**. Representative images of regions of healthy and bleomycin treated mouse lung stained with fast green FCF. **B**. Quantification of fast green FCF staining as mean intensity of fluorescence signal in lung parenchyma. **C**. Quantification of fast green FCF staining as percentage of lung parenchyma stained. **D**. Image of paraffin-embedded sections from the same samples stained with Masson’s trichrome, a stain that clearly labels collagen in blue but is limited by morphology and sensitivity of bright-field thin section imaging. **E**. Samples stained with picrosirius red, which stains collagen in red but shows non-specific staining of cytoplasm in mixed inflammatory/fibrotic lesions after bleomycin treatment (black arrowhead). **F**. Specificity is not improved when picrosirius red-stained samples are viewed using fluorescence microscopy (yellow arrowhead). **B & C** show means ± SEM, n=5 per group, *P*-values from an unpaired t-test on log_10_-transformed data.

**Figure 3.**
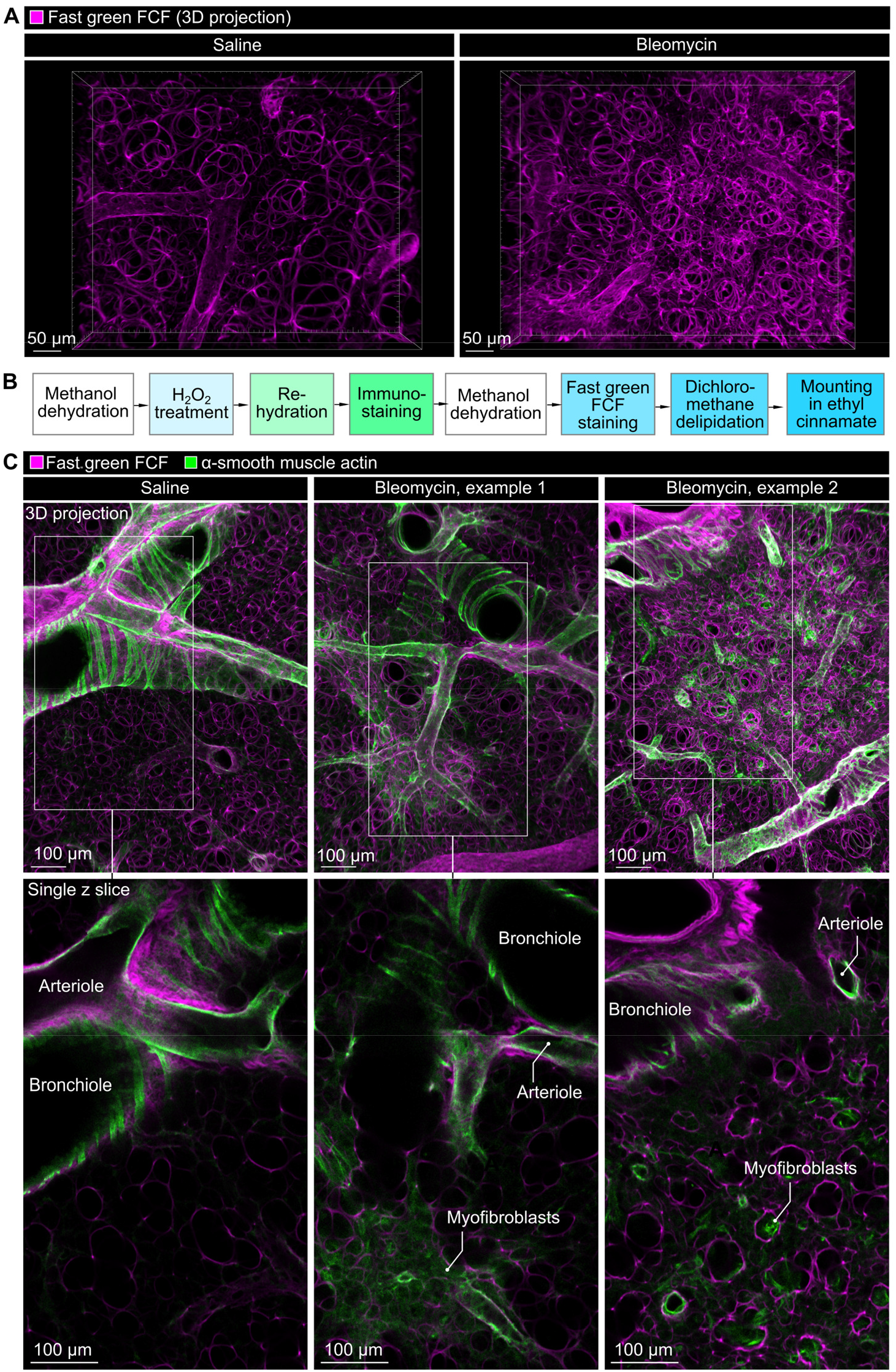
Fast green FCF staining enables 3D imaging of fibrotic lesions and can be imaged in combination with immunofluorescence. **A**. 3D imaging of control and bleomycin-treated mouse lung parenchyma (88 µm thickness, 2 µm z increments). **B**. Workflow for combining immunostaining with fast green FCF staining. **C**. Fast green FCF staining combined with immunostaining for α-smooth muscle actin (α-SMA) in mouse lung slices, showing 3D projections (72 µm thickness, 2 µm z increments, upper images) and single z slices (lower images). Images are representative of 5 samples.

Fluorescence approaches have the advantage over brightfield staining of allowing simultaneous imaging with immunofluorescence stains. Staining for α-SMA provides useful structural information, can reveal myofibroblast proliferation and can be used to measure pathological arterial muscularization when studying pulmonary hypertension related to interstitial lung disease. We therefore developed an approach for combining fast green FCF staining with immunofluorescence staining for alpha-smooth muscle actin (α-SMA) (**Fig 3A**), which successfully enabled simultaneous imaging of α-SMA and collagen in 3D (**Fig 3B**).

Another advantage of cleared tissue samples stained with fluorescent dyes is that these preparations are compatible with light sheet fluorescence microscopy, which enables greatly accelerated volumetric imaging relative to point-scanning approaches. Oblique plane microscopy is a light sheet modality that is well-suited to higher-throughput quantitative imaging in preclinical assays as it is compatible with ‘open-top’ microscopes capable of scanning organ slice or organoid samples in multi-well plates^14,15^. Oblique plane scans enabled rapid 3D mapping of collagen networks in mouse lung slices (**Fig 4A**).

**Figure 4.**
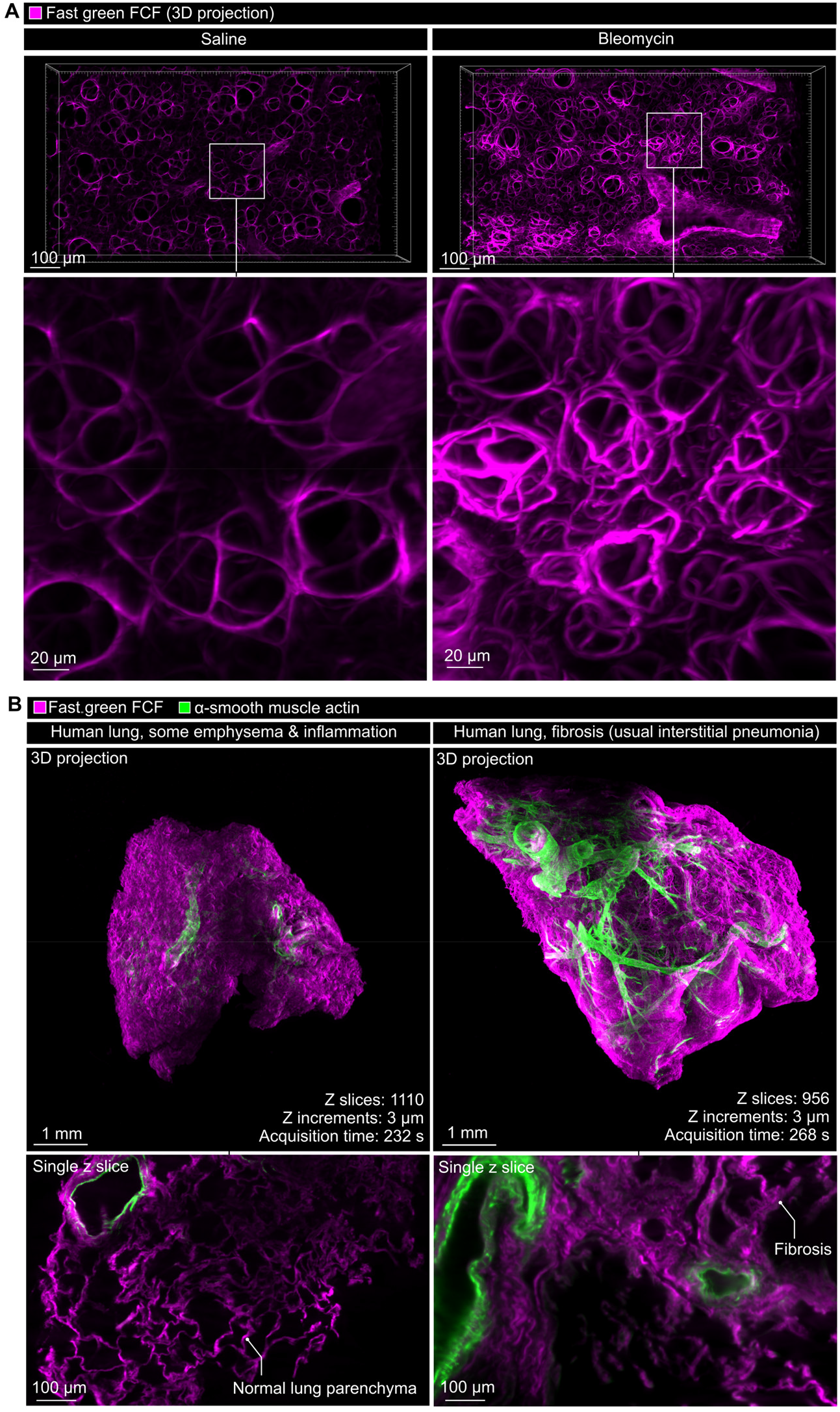
Fast green FCF-staining enables rapid 3D imaging of mouse and human lung samples using light sheet fluorescence microscopes. **A**. Images of saline control and bleomycin-treated mouse precision-cut lung slices scanned using oblique plane microscopy (0.1 mm^3^ sampled, 0.5 µm z sampling interval after deskewing, total acquisition time of 340 s per 3D volume for a single fluorescence channel without acceleration that could be achieved with hardware triggering). **B**. Images of human lung samples from background lung obtained during cancer resections from a patient with no fibrotic changes observed (left) and from a sample where fibrosis was noted in clinical pathology workup (usual interstitial pneumonia pattern, right) (6.4 mm^3^ sampled, 3 µm z increments, 268 seconds per image for two colour channels).

A current goal in clinical pathology is to achieve entire-biopsy 3D microscopy scans to overcome the fundamental problem of under-sampling when only thin sections are imaged^16^. This under-sampling problem is relevant to pulmonary fibrosis research as fibrotic pathology is typically patchy in its presentation and early changes can be hard to detect^17^. We therefore stained and cleared human lung samples with and without pathologist-determined fibrosis in preparations suitable for whole-sample light sheet fluorescence microscopy scanning. This approach also enabled rapid 3D imaging of collagen in combination with α-SMA immunofluorescence staining in human lung samples (**Fig 4B**).

Together, our findings demonstrate that fast green FCF staining is a useful tool for imaging fibrosis in preclinical assays and may also have use for achieving 3D clinical pathology.

## Discussion

In this study we have confirmed that fast green FCF staining is useful for imaging pulmonary fibrosis. Fast green FCF staining has several advantages over histology stains that are in current widespread use, which include the high specificity of fast green FCF for fibrillar collagen, its compatibility with 3D imaging, and its suitability for quantifying collagen from fluorescence intensity. Fast green FCF is also less expensive and exhibits improved tissue penetration relative to antibodies used for immunofluorescence imaging of type I collagen^7^.

A notable limitation of fast green FCF is that this dye requires staining and imaging in anhydrous conditions for bright collagen labelling. In our work we have moved away from using dibenzyl ether, an irritant and environmentally hazardous mounting media used in previous studies^7,11^, towards using lower-toxicity ethyl cinnamate^18^. In future it may be also useful to identify strategies to avoid use of methanol and dichloromethane as these solvents require handling with care due to toxicity and incompatibility with many types of plastic^18^. We also noted uneven signal intensity adjacent to collagen-rich regions in larger human tissue preparations. This appears to be due to attenuation of fluorescence signal by fast green FCF dye in the light sheet path and in ongoing work we are assessing whether staining with lower concentrations of fast green FCF improves images of fibrotic samples with a thickness of several millimeters.

In addition to the animal model and clinical samples imaged in this study, it would be of interest to test whether the high sensitivity of fast green FCF staining enables measurement of collagen remodeling in human precision-cut lung slices treated with fibrotic cocktail mediators. These ex vivo assays are increasingly used to study fibrogenesis in vitro, but have a need for protein-level readouts that are both quantifiable and have clear links to fibrosis^19,20^. The protocol we developed for imaging mouse precision-cut lung slices provides a starting point for this work. Fast green FCF staining approaches may also be useful to study fibrosis occurring in other organs. Because fast green FCF is bright, specific and amenable to 3D imaging of dense tissue, we anticipate that this dye will become a widely-used tool for imaging fibrosis in the lungs and beyond.

## Methods

### Animals

C57BL/6 mice (C57BL/6NCrl, Charles River) were bred and maintained in specific pathogen-free conditions at King’s College London. Male and female mice aged 8-12 weeks were used for experiments. Animals were randomly allocated to treatment groups within cage groups. Procedures were authorized under license PP1060882.

### Bleomycin-induced pulmonary fibrosis model

Mice were anaesthetised with isoflurane and given a single dose of 2000 IU/kg bleomycin (Accord) or vehicle control (normal saline) at 2 µl/g body weight by oropharyngeal aspiration^21^.

### Mouse lung processing

At 2 weeks post saline/bleomycin administration, mice were anesthetised with isoflurane, exsanguinated and perfused via the right ventricle with 20 ug/ml heparin in PBS for 30 seconds and then 1% formaldehyde in PBS for 2 minutes at 25 mmHg. Lungs were then inflated with 40 µl/g body weight of molten 2% agarose in PBS. Agarose was allowed to set before lungs were excised, washed in PBS and fixed overnight in 4% formaldehyde in PBS at 4 °C. Precision-cut lung slices were produced at 250 µm thickness using a vibratome (Precisionary VF-510-0Z).

### Human lung processing

Background lung samples, collected at regions distal from tumor margins during cancer resections, were obtained from the Heart, Lung and Critical Care Biobank, Royal Brompton & Harefield Hospitals, part of Guy’s and St Thomas’ NHS Foundation Trust (research ethics committee review code: 08/H0407). Surgical lung samples up to 4 cm in diameter were sub-sampled to produce samples the size of transbronchial biopsies^22^.

### Fast green FCF staining, immunofluorescence and tissue clearing

Staining approaches for different preparations are detailed in **Table S1**. Adaptations to previous approaches include use of ethyl cinnamate as mounting media, which is less toxic than dibenzyl ether^18^, and use of CUBIC-L for enhancing clearing of larger, non-perfused fibrotic lung samples, which is more effective than dichloromethane at eluting heme from residual blood^8,23,24^. Precision-cut lung slices were mounted in ethyl cinnamate in a 24 well glass bottom plate (Ibidi 82427), held in position using a weighted coverslip (**Fig S1A**). To facilitate handling, human lung samples were embedded in 2% low-melting agarose in deionized water before dehydration, fast green FCF staining and refractive index matching imaging immersed in ethyl cinnamate (**Fig S1B**).

For immunostaining, samples were stained with anti-α-SMA-FITC (Sigma-Aldrich F3777) at 1:1000 (2.7 µg/ml) in 300 ul per lung slice in PBS + 0.3% triton X-100 + 0.5% normal donkey serum. The same antibody was used to stain human lung samples, at 1:250 (10.8 µg/ml) in 2 ml per sample HEPES-TSC buffer (200 mM NaCl, 10 mM HEPES, 10% triton X-100, 0.05% sodium azide, 0.5% casein, pH 7.5) for 7 days at 37°C^24,25^. After washing, bound antibodies were fixed in position with 1% formaldehyde before proceeding with dehydration for fast green FCF staining.

### Microscopy

Mouse lung slices were imaged with a Nikon AXR with NSPARC using a 10X objective (Nikon Imaging Centre at King’s). Excitation/emission data were gathered using the lambda scan function on a Leica Stellaris 8 system (Microscopy Innovation Centre at King’s). Conventional histology preparations were imaged with an Olympus BX51 using a 20X objective.

Oblique plane light sheet microscopy was achieved using a custom system based on the original design of this modality^15^, built around a Nikon Eclipse Ti2-E system at the Microscopy Innovation Centre at King’s. This system used an imaging objective (Nikon Plan Apo λD 60X/1.42 Oil) as well as two remote-focusing objectives (Nikon Plan Apo λ 40X/0.95 and Nikon Plan Fluor ELWD 40X/0.60, respectively) for re-imaging and detection. For oblique plane microscopy, fast green FCF was excited at 561 nm, with excitation separated from fluorescence emission using a quad-band dichroic mirror (DC/ZT405/488/561/640rpcv2-UF3, Chroma Technology). Emitted fluorescence was filtered using a Semrock 617/73 nm bandpass filter and detected with a Hamamatsu ORCA-Flash4.0 sCMOS camera. Human lung samples were imaged using a commercially available UltraMicroscope Blaze light sheet system with a 4X objective (Miltenyi Biotec).

### Image analysis

Measurements were made from images using QuPath 0.7.0. Analysis was restricted to lung parenchyma, with signal from extrapulmonary structures and cuff spaces surrounding large bronchovascular bundles and pulmonary veins excluded as they contain large quantitates of fibrillar collagen at baseline. 3D projections were rendered using Imaris 11.

### Histology

After collection of vibratome sections, lung blocks were dehydrated and processed for embedding in paraffin wax blocks. Sections (4 µm thickness) were sampled from the block immediately adjacent to regions sampled for fast green FCF staining. Slides were stained using standard Masson’s trichrome (omitting nuclear stain) and picrosirius red staining protocols^13,26^.

### Statistics

Data are expressed as mean ± standard error. Statistical testing was conducted as described in figure legends, using GraphPad Prism 11.0.2, with P-values <0.05 considered statistically significant.

## Data availability

Datasets shown in graphs are provided in the **Data Supplement**. Additional detail on timings and volumes and approaches for preparations are provided in the supplemental files (**Table S1, Figure S1**). Image files are available on request.

## Acknowledgements

SJC is grateful for funding from an Academy of Medical Sciences Springboard award and the Bioimaging UK user access fund. MS was supported by a British Pharmacological Society vacation studentship. AKR was supported by a British Association for Lung Research-Action for Pulmonary Fibrosis summer studentship. We thank the Nikon Imaging Centre and Microscopy Innovation Centre at Kings College London for support with light microscopy, with special thanks to Vincenzo Infante. Human lung imaging was possible thanks to Jak Grimes and Nimo Abdullah (Miltenyi Biotec), Alessandro Ciccarelli (Francis Crick Institute) as well as support from the Heart, Lung & Critical Care biobank team and the generosity of patients who donated lung tissue.

## Author contributions

Conceptualization: SJC; Data curation: MS, AKR, DH, SJC; Formal analysis: MS, AKR, SJC; Funding acquisition: SJC; Investigation: MS, AKR, SM, JRB, AB, AAJ, DH, CH, SJC; Methodology: SJC, AB, AAJ, DH, CH; Resources: JRB, SJC; Supervision: SJC; Writing – original draft: SJC; Writing – review and editing: all authors.

## Saqib & Rivers et al. - supplemental tables and figures

**Table S1.**
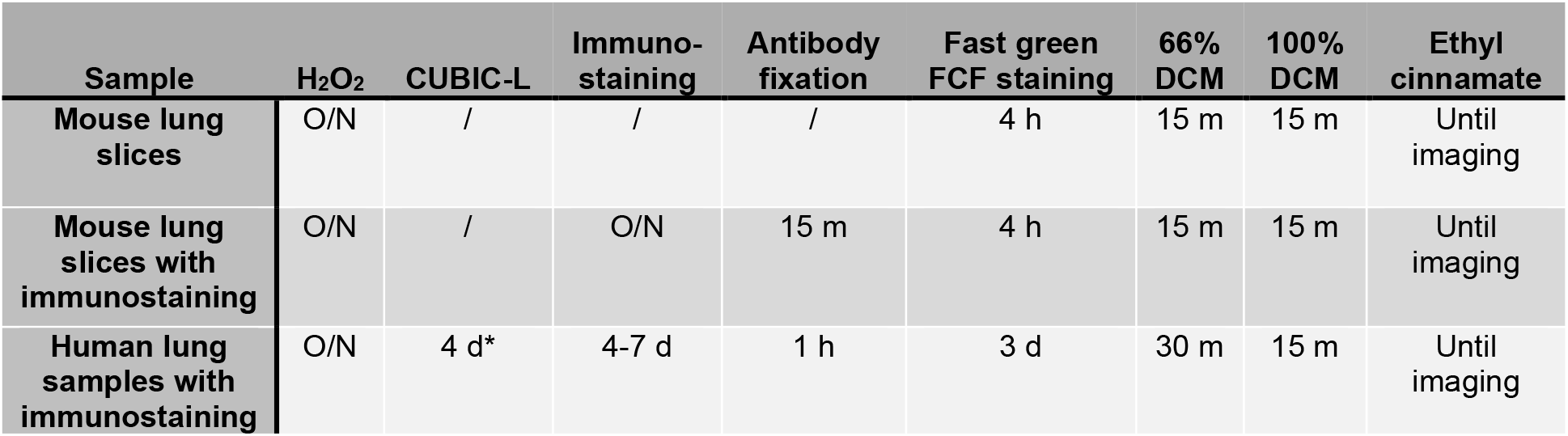
Summary of processing steps and incubation times for tissue clearing and staining. Abbreviations: O/N = overnight; / = not performed step; DCM = dichloromethane. Details on dehydration, rehydration and wash steps are omitted for brevity but were performed as described in the **Methods** section and as shown in **Fig 1A** and **Fig 3B**^1^. For other steps, volumes used for mouse lung slices were: 300 µl (immunostaining) and 500 µl (washes) and for human lung samples: 2 ml (CUBIC-L treatment, immunostaining) and 3 ml (washes).

| Sample | H<sub>2</sub>O<sub>2</sub> | CUBIC-L | Immuno-staining | Antibody fixation | Fast green FCF staining | 66% DCM | 100% DCM | Ethyl cinnamate |
| --- | --- | --- | --- | --- | --- | --- | --- | --- |
| Mouse lung slices | O/N | / | / | / | 4 h | 15 m | 15 m | Until imaging |
| Mouse lung slices with immunostaining | O/N | / | O/N | 15 m | 4 h | 15 m | 15 m | Until imaging |
| Human lung samples with immunostaining | O/N | 4 d* | 4-7 d | 1 h | 3 d | 30 m | 15 m | Until imaging |

**Figure S1.**
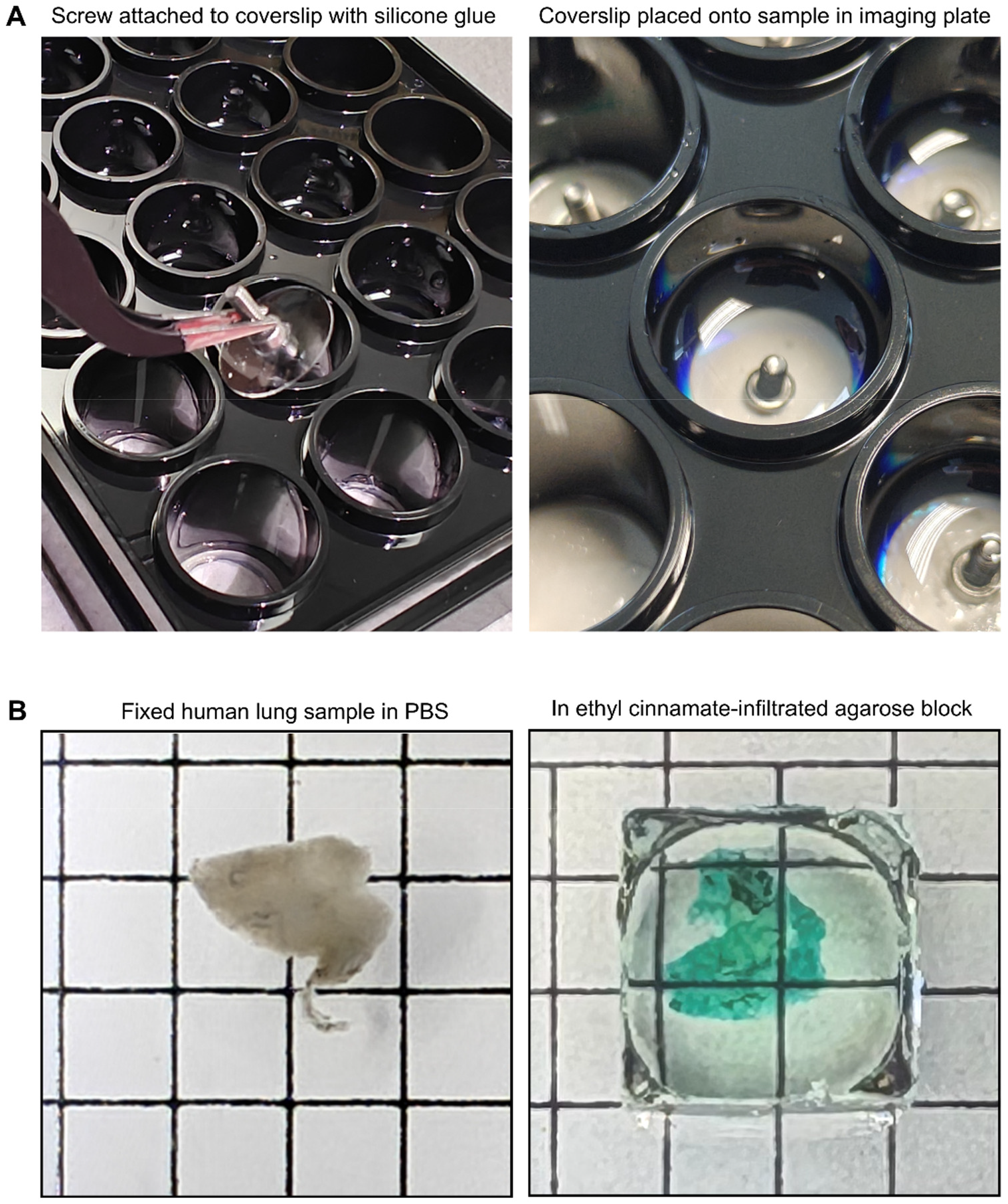
Techniques for mounting cleared samples in ethyl cinnamate. **A**. Glass bottom 24-well plates (Ibidi, 82427) can be used with ethyl cinnamate. To hold lung slices flat and for securing them during tiled scans, a 13 mm diameter coverslip is placed on top of the lung slice. Screws were attached to the coverslip as shown, with silicone glue (Henkel Loctite 595), for handling. Warning: although secure for over a week, after several weeks of exposure to ethyl cinnamate the glue attaching the coverglass to the 24-well plate will be weakened and can leak. Ethyl cinnamate degrades the plate lid so wells can be covered with parafilm. Plates containing ethyl cinnamate can be stored inside containers made from polypropylene. **B**. To facilitate handling and mounting for light sheet microscopy, human lung samples were embedded in agarose before dehydration, fast green FCF staining and refractive index matching.

## References

1. Rajan, S. K. et al. Progressive pulmonary fibrosis: an expert group consensus statement. European Respiratory Journal 61, (2023).

2. Raghu, G. et al. Diagnosis of Idiopathic Pulmonary Fibrosis. An Official ATS/ERS/JRS/ALAT Clinical Practice Guideline. Am J Respir Crit Care Med 198, e44–e68 (2018).

3. Fabbri, L. et al. Research priorities for progressive pulmonary fibrosis in the UK. BMJ Open Resp Res 11, (2024).

4. Jenkins, R. G. et al. An Official American Thoracic Society Workshop Report: Use of Animal Models for the Preclinical Assessment of Potential Therapies for Pulmonary Fibrosis. Am J Respir Cell Mol Biol 56, 667–679 (2017).

5. Ashcroft, T., Simpson, J. M. & Timbrell, V. Simple method of estimating severity of pulmonary fibrosis on a numerical scale. J Clin Pathol 41, 467–470 (1988).

6. Baluk, P. et al. Lymphatic Proliferation Ameliorates Pulmonary Fibrosis after Lung Injury. The American Journal of Pathology 190, 2355–2375 (2020).

7. Timin, G. & Milinkovitch, M. C. High-resolution confocal and light-sheet imaging of collagen 3D network architecture in very large samples. iScience 26, (2023).

8. Tsukui, T., Wolters, P. J. & Sheppard, D. Alveolar fibroblast lineage orchestrates lung inflammation and fibrosis. Nature 631, 627–634 (2024).

9. Belmans, F. et al. Imaging technologies in experimental pulmonary fibrosis research: essential tool for enhanced translational relevance. European Respiratory Review 34, (2025).

10. Song, S. et al. Intracellular hydroxyproline imprinting following resolution of bleomycin-induced pulmonary fibrosis. European Respiratory Journal 59, (2022).

11. Pechtimaldjian, L. et al. Skin-iDISCO+: An optimized tissue-clearing and labeling protocol for morphometric analysis of human cutaneous vasculature. STAR Protoc 6, 103524 (2025).

12. West, J. B. Comparative Physiology of the Pulmonary Circulation. Comprehensive Physiology 1, 1525–1539 (2011).

13. Junqueira, L. C., Bignolas, G. & Brentani, R. R. Picrosirius staining plus polarization microscopy, a specific method for collagen detection in tissue sections. Histochem J 11, 447–455 (1979).

14. Sparks, H. et al. High content 3D imaging by dual-view oblique plane microscopy. PNAS Nexus 4, pgaf370 (2025).

15. Dunsby, C. Optically sectioned imaging by oblique plane microscopy. Opt. Express, OE 16, 20306–20316 (2008).

16. Forjaz, A. et al. Three-dimensional assessments are necessary to determine the true, spatially resolved composition of tissues. Cell Rep Methods 5, 101075 (2025).

17. Wolters, P. J., Collard, H. R. & Jones, K. D. Pathogenesis of Idiopathic Pulmonary Fibrosis. Annu Rev Pathol 9, 157–179 (2014).

18. Masselink, W. et al. Broad applicability of a streamlined ethyl cinnamate-based clearing procedure. Development 146, dev166884 (2019).

19. Alsafadi, H. N. et al. An ex vivo model to induce early fibrosis-like changes in human precision-cut lung slices. American Journal of Physiology-Lung Cellular and Molecular Physiology 312, L896–L902 (2017).

20. Lehmann, M. et al. Precision-Cut Lung Slices: Emerging Tools for Preclinical and Translational Lung Research: An Official American Thoracic Society Workshop Report. Am J Respir Cell Mol Biol 72, 16–31 (2025).

21. Seo, Y. et al. Optimizing anesthesia and delivery approaches for dosing into lungs of mice. Am J Physiol Lung Cell Mol Physiol 325, (2023).

22. Carducci, C., Kaminski, N., Justet, A. & Tomassetti, S. Cryobiopsy at the dawn of fibrosis: unlocking early diagnosis and therapeutic windows. Front. Med. 13, (2026).

23. Susaki, E. A. et al. Whole-brain imaging with single-cell resolution using chemical cocktails and computational analysis. Cell 157, 726–739 (2014).

24. Cleary, S. J. et al. Intravital imaging of pulmonary lymphatics in inflammation and metastatic cancer. J Exp Med 222, e20241359 (2025).

25. Susaki, E. A. et al. Versatile whole-organ/body staining and imaging based on electrolyte-gel properties of biological tissues. Nat Commun 11, 1982 (2020).

26. Dowland, S. Masson’s Trichrome staining for histology. protocols.io 10.17504/protocols.io.e6nvwbwwwvmk/v1 (2025).

